# Genetic drift opposes mutualism during spatial population expansion

**DOI:** 10.1101/003012

**Authors:** Melanie J. I. Müller, Beverly I. Neugeboren, David R. Nelson, Andrew W. Murray

## Abstract

Mutualistic interactions benefit both partners, promoting coexistence and genetic diversity. Spatial structure can promote cooperation, but spatial expansions may also make it hard for mutualistic partners to stay together, since genetic drift at the expansion front creates regions of low genetic and species diversity. To explore the antagonism between mutualism and genetic drift, we grew cross-feeding strains of the budding yeast S. cerevisiae on agar surfaces as a model for mutualists undergoing spatial expansions. By supplying varying amounts of the exchanged nutrients, we tuned strength and symmetry of the mutualistic interaction. Strong mutualism suppresses genetic demixing during spatial expansions and thereby maintains diversity, but weak or asymmetric mutualism is overwhelmed by genetic drift even when mutualism is still beneficial, slowing growth and reducing diversity. Theoretical modeling using experimentally measured parameters predicts the size of demixed regions and how strong mutualism must be to survive a spatial expansion.

## 1 Introduction

Spatial population expansions are common events in evolutionary history. They range from the growth of microbial biofilms on surfaces [1] to the pre-historic human migration out of Africa [2] and will occur more frequently as climate change forces species to shift their territories [3]. When populations expand, the first individuals to arrive in the new territory are likely to be the ancestors of the later populations in this area. This ‘founder effect’ produces regions with low genetic diversity because they are occupied by the progeny of a few founders [4]. With few founders, the random sampling of individuals (genetic drift) becomes important. The invasion of different regions by different founders can lead to spatial separation of genotypes (‘demixing’) [4, 5].

Territorial expansions can have profound effects on the interactions between species or genotypes [6, 7]. For example, the associated demixing can spatially separate cooperators from non-cooperating ‘cheaters’ [8, 9, 10, 11], in line with the common view that spatial structure in general enhances cooperation. In contrast, spatial demixing may have a detrimental effect on mutualistic interactions (beneficial for both partners). Mutualism selects for coexistence (‘mixing’) of the two partners [12], as was recently shown for a microbial mutualism in a spatial setting [13], and theory argues that the demixing caused by spatial expansion can extinguish mutualism [14, 15]. Mutualism imposes constraints on spatial expansions: Obligate mutualists must invade new territory together, and facultative mutualists invade faster when mixed.

Despite these constraints, major events in evolutionary history involve spatial expansions of mutualists. The invasion of land by plants may have taken advantage of the mutualistic association with fungi [16], and flowering plants spread with their pollen-dispersing insects [17]. More recently the invasion of pine trees in the Southern hemisphere required mycorrhizal fungal symbionts [18], and legumes can only grow in new areas with their mutualist nitrogen-fixing rhizobacteria [19]. Microbes in biofilms often exhibit cooperative interactions [20], such as interspecies cooperation during tooth colonization [21]. A common microbial mutualism is cross-feeding, i.e. the exchange of nutrients between species [22, 23, 24, 25, 26, 27].

Here, we use the growth of two cross-feeding strains of the budding yeast *Saccharomyces cerevisiae* on agar surfaces to study the antagonism between genetic drift and mutualism during spatial expansions. The strains exchange amino acids, allowing us to control the mutualism’s strength by varying the amino acid concentrations in the medium. The strains demix under non-mutualistic conditions, but, for obligate mutualism, expand in a more mixed pattern whose characteristics we explain with a model of the nutrient exchange dynamics. When mutualism is facultative or highly asymmetric, genetic drift dominates, leading to demixing even when mixing would be beneficial. We quantitatively understand this transition using a generalized stochastic Fisher equation.

**Figure 1:**
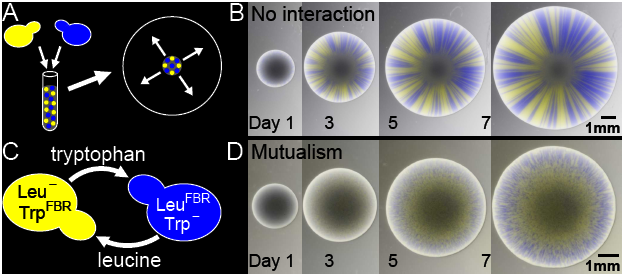
**A)** Spatial expansion assay. Two fluorescently labeled *S. cerevisiae* strains, depicted as blue and yellow, are mixed in liquid and pipetted as a circular drop onto an agar surface. When the colony expands, the ensuing spatial pattern can be monitored by fluorescence microscopy. **B)** Successive images of the expansion of two non-interacting yeast strains show the formation of distinctive blue and yellow sectors. **C)** Two cross-feeding yeast strains as a model for mutualism. The yellow strain Leu^FBR^ Trp^−^ produces leucine but not tryptophan, while the blue strain Leu^−^ Trp^FBR^ produces tryptophan but not leucine. To grow on medium lacking both amino acids, the strains must cross-feed each other. The strains are feedback-resistant (FBR) in the production of leucine or tryptophan, leading to increased production and therefore secretion of these amino acids. **D)** These mutualistic strains form small, intertwined patches during spatial expansion, see also Fig. 2C.

## 2 Results

To study spatial expansions, Hallatschek *et al.* pioneered a simple microbial expansion assay [5]. Two yeast strains labeled with two different fluorescent proteins, depicted as yellow and blue in Fig. 1A, are mixed and inoculated as a circular drop (the ‘homeland’) on an agar surface. The colony grows radially outwards on the surface as cell division pushes cells forward (yeast has no active motility). The cells deplete the nutrients in the agar immediately below the colony, and then grow solely on nutrients diffusing towards the colony from the surrounding agar, restricting growth to a small ‘active layer’ extending only 40 *μm* back from from the colony boundary [28, 29]. The small number of cells involved in local colony propagation leads to a high local fixation probability for blue or yellow cells [5, 30] (Fig. 1B). Colony expansion reduces diversity: a front that migrates from a well-mixed homeland produces sectors that are fixed for yellow or blue cells.

### 2.1 Strong mutualism inhibits demixing

To study mutualism, we genetically engineered the yeast strains shown in Fig. 1C. These strains cross-feed each other two amino acids, leucine (leu) and tryptophan (trp). To enhance cross-feeding, we used previously characterized feedback-resistant (FBR) mutations [31, 32] that increase amino acid production by inactivating the feedback inhibition that normally regulates amino acid production. The strain Leu^FBR^ Trp^−^, depicted as yellow, overproduces leucine (Leu^FBR^) and leaks it into the medium, but cannot produce tryptophan (Trp^−^). Its partner strain Leu^−^ Trp^FBR^, depicted as blue, overproduces and leaks tryptophan (Trp^FBR^) but cannot produce leucine (Leu^−^). Because growth requires both leucine and tryptophan, neither strain can grow on medium lacking both amino acids.

When mixed together, the two cross-feeding strains grow robustly on medium without leucine and tryptophan, forming interdigitated patches (Fig. 1D). Since each strain needs the amino acid from its partner strain to proliferate, they cannot demix into the large separated sectors seen for non-interacting strains, but must stay in close proximity. However, some segregation still occurs: the mutualists make visible yellow and blue patches (Fig. 1D), but these patches are much smaller than the sectors of non-interacting strains (Fig. 1B).

**Figure 2:**
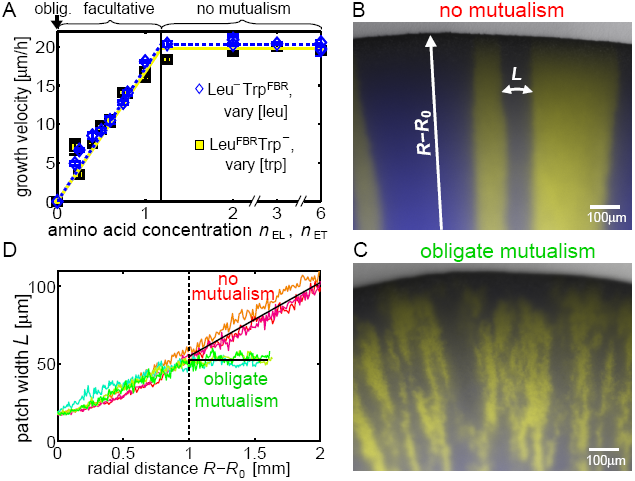
**A)** The radial growth velocities of single-strain colonies of Leu^−^ Trp^FBR^ (blue diamonds) and of Leu^FBR^ Trp^−^ (yellow squares) increase linearly with the leucine concentration *n*_EL_ = [leu]/[leu]_c_ and the tryptophan concentration *n*_ET_ = [trp]/[trp]_c_ in the medium, respectively, until they saturate at a plateau. This behavior is indicated by the corresponding blue dotted and yellow solid lines, which are piecewise linear fits. Concentrations are scaled with the factors [leu]_c_ = 762 *μ*M and [trp]_c_ = 98 *μ*M, to make the crossover between the linear and the plateau regime occur at the same rescaled concentrations (vertical line). For concentrations above this crossover value, mutualism is irrelevant *(‘no mutualism*’). Mutualism is *facultative* for lower and *obligate* for zero concentrations. **B)** For no mutualism, colonies exhibit demixing into large sectors. **C)** For obligate mutualism, colonies form much smaller, intertwined patches. **D)** Average width *L* (parallel to the front) of yellow and blue patches as function of the radial distance *R* – *R*_0_ from the homeland (perpendicular to the front) for three replicate colonies under conditions of no (three independent, reddish lines) and obligate (three independent, greenish lines) mutualism. For the first mm, *L* increases due to genetic demixing and sector boundary diffusion. Afterwards, obligate mutualism limits the patch width to *L* = 52 *μ*m (black horizontal line), while the sector width increases linearly with the radius for no mutualism (black inclined line).

Genetic drift and mutualism are opposing forces: mutualism mixes and drift demixes. To probe this antagonism, we change the strength of mutualism by varying the levels of leucine and tryptophan in the medium. We quantify this effect by measuring the expansion velocities of single-strain colonies for different concentrations of the amino acid they need, e.g. leucine for Leu^−^ Trp^FBR^(Fig. 2A). As expected, strains cannot grow without their required amino acid; as the amino acid concentration increases, the velocity increases approximately linearly and then plateaus. The surprising linearity below the plateau suggests that cells alter the number and/or the affinity of amino acid transporters in response to the external amino acid concentration. Appropriately scaling the amino acid concentrations makes velocity plots of the two strains overlap almost exactly. The leucine scaling factor, 762 *μ*M, is 8 times larger than the tryptophan scaling factor, 98 *μ*M;, consistent with yeast proteins containing ~10 times as much leucine as tryptophan [33]. The small deviation of the vertical crossover line from a value of 1 in Fig. 2A is due to amino acid loss into the medium as explained in SI Sec. S2.

The effects of amino acid concentration on growth define three growth regimes. On medium without leucine and tryptophan, the two strains can grow together but not alone and thus form a pair of *obligate mutualists.* For increased amounts of leucine and tryptophan, mutualism becomes *facultative,* since the strains get amino acids from the medium as well as from their partner (that the partner’s presence still leads to faster growth in this regime is shown below in Fig. 4F). Above a critical concentration (the onset of the plateau regime of Fig. 2A), leucine and tryptophan are no longer growth limiting, and amino acids leaked by the cells should not matter. We therefore expect cells to demix into well-defined sectors like the non-interacting cells in Fig. 1B. This is indeed the case (Fig. 2B), defining this as the *no mutualism* regime.

### 2.2 Patterns for obligate and no mutualism

We first studied the extreme cases of obligate and no mutualism with their striking difference in expansion patterns (Fig. 2B,C). To quantify this difference, we used image analysis to determine how the average width *L* (parallel to the front) of patches of a single color changes as expansion progresses for increasing radial distances *R* — *R*_0_ from a homeland of radius *R_0_.* As shown in Fig. 2D, for both obligate and no mutualism the patch width initially increases as unicolored patches form by local fixation events due to genetic drift. In addition, patch boundaries diffuse and create larger and larger patches when they collide [5, 34]. For radii *R* larger than twice the homeland radius *R*_0_ ≈ 1 mm, the no-mutualism sector width increases linearly with the radius because the sector width increases with the growing colony’s circumference, preventing further sector boundary collisions so that the sector number stays constant [34].

For obligate mutualism the patch width plateaus at *L* = 52 ± 1*µ*m, even for large radii. Mutualism requires physical proximity of the interacting partners. For example, cells in a patch of leucine-requiring cells get leucine by diffusion from neighboring leucine-producing patches. The patch cannot get too big. If it did, the cells at its center would starve, because the more peripheral cells would take up all the leucine diffusing from the neighboring patches of leucine producers.

We set out to combine theory and experimentally measured parameters to understand the patch width and other characteristics of the expansion pattern from the nutrient exchange dynamics. The dynamics of the leucine concentration [leu](*x,t*) in a reference frame that moves with the front is

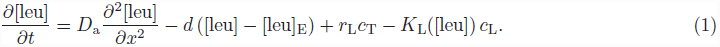

The first term describes leucine diffusion, with diffusion constant *D*_a_, along the coordinate *x* parallel to the front. The second, chemostat-like term is an effective description of leucine diffusion perpendicular to the front. The gradient between the leucine concentration [leu] at the colony boundary and the leucine concentration [leu]_E_ in the medium far away from the colony generates a diffusive flux away from the colony with the diffusive rate *d* (see SI Sec. S5 for a detailed derivation). The third term describes secretion of leucine by leucine-producing cells, whose concentration is *c*_T_, with rate *r*_L_ per cell, while the last term describes leucine uptake by leucine-requiring cells, whose concentration is *c*_L_, with a concentration-dependent rate *K*_L_([leu]). To maintain constant intracellular amino acid concentrations during steady-state growth, the leucine uptake rate has to be proportional to the growth rate. The dependence of colony growth velocity on external amino acid concentration (Fig. 2A) thus motivates a constant uptake rate *K_L_*([leu]) = *k_L_* for concentrations larger than the crossover value [leu]_c_, and linear *K*_L_([leu]) = *k*_L_[leu]/[leu]_c_ for smaller concentrations. Such a limiting behavior for small and large concentrations is also expected for other functional forms of the uptake rate such as Michaelis-Menten kinetics.

We write Equation [1] in a non-dimensionalized form for the rescaled concentration *n*_L_ ≡ [leu]/[leu]_c_ by expressing time in units of the inverse diffusive rate *d* and space in units of the diffusion length scale: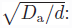

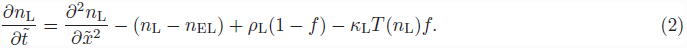

Here, we have written the cell concentrations C_L_ and C_T_ in terms of the fraction *f≡ C*_L_/(*C*_L_ + *C*_T_) of leucine-requiring cells, where *C* = *C*_L_ + *C*_T_ is the constant surface carrying capacity (yeast colonies grow to a finite height). We have defined the function *T*(*n*_L_) = *n*_L_ for *n*_L_ ≤ 1 and = 1 for *n*_L_ > 1, and the dimensionless secretion and uptake parameters

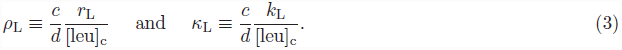

Similar equations hold for tryptophan with the rescaled secretion and uptake rates *ρ*_T_ ≡ (*c/d*)(*r*_T_/[trp]_c_) and *κ*_T_ ≡ (*c/d*)(*k*_T_/[trp]_c_). We find that these parameters are equal for our two strains, *ρ*_L_ = *ρ*_T_ ≡ *ρ* and *κ*_L_ = *κ*_T_ ≡*κ* (SI Sec. S1.4). Thus, the mutualism described by Equation [2] and the corresponding tryptophan equation is a symmetric interaction, unless asymmetries are introduced via the external concentrations *n*_EL_ and *n*_ET_. This symmetry is not trivial (below we will see that it does not hold away from steady-state growth), but presumably not accidental. Since yeast cells contain an order of magnitude more leucine than tryptophan, we expect uptake and secretion to be an order of magnitude larger for leucine than for tryptophan. Indeed, the equalities *ρ*_L_ = *ρ*_T_ and *κ*_L_ = *κ*_T_ follow from the scaling relations *k*_L_ = 8 *k*_T_, *r_L_* = 8*r*_T_, and [leu]_c_ = 8 [trp]_c_ in our system (SI Sec. S1.4).

According to the symmetry for obligate mutualists (*n*_EL_ = *n*_ET_ = 0), blue and yellow patches should have the same average widths, which we observe (Fig. 2C and SI Fig. S4A). Although mutualist patches are less clearly defined than demixed non-mutualist sectors, we first assume, for simplicity, that a blue patch contains only leucine consumers (*f* = 1). Leucine diffuses into such a patch from neighboring yellow leucine producer patches, but is lost due to uptake and diffusion away from the colony with the combined rate *ck*_L_/[leu]_c_ + *d* = *dκ* + *d*. On the loss time scale *t*_loss_, which equals the inverse loss rate, leucine diffuses a distance 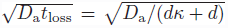 into the consumer patch. More rigorously, according to Equation [2] the leucine concentration within the patch decreases exponentially with the distance from the neighboring leucine-producing patches on the length scale

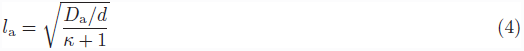

This gradient would lead to the patch growing more slowly in its middle than at its boundary, which would cause an unstable, undulating front rather than the smooth front seen in Fig. 2C. Thus, patches must be small compared to the length scale *l*_a_ that describes the fall of nutrient concentration within a patch. Indeed, for our system *l*_a_ ≈ 700 *μ*m, an order of magnitude larger than the mutualistic patch width *L* ≈ 50 *μ*m.

Patch boundaries wander perpendicular to the expansion direction due to the jostling of cell division [34]. Boundary diffusion can ‘smooth out’ velocity differentials caused by amino acid diffusion provided that both processes happen on comparable length and time scales. A quantitative calculation (SI Sec. S4) shows that this is the case for an average patch width

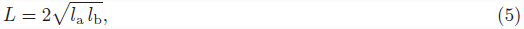

which is twice the geometric mean of the nutrient diffusion length scale *l*_a_ and the patch boundary diffusion length *l*_b_ = 2*D*_s_/*b*. Since the cellular diffusion constant *D*_s_(a few *μ*m^2^/h) is much smaller than the amino acid diffusion constant *D*_a_(a few mm^2^/h), the length scale for the diffusion of the boundary between the two cell types in the active layer of size *b* is only *l_b_* ≈ 1 *μ*m compared to *l_a_* ≈ 700 *μ*m. From Equation [5], we estimate *L* ≈ 50 *μ*m, which agrees well with the value observed in Fig. 2.

Due to patch boundary diffusion, patches have ragged boundaries and are not completely demixed. Patches that look yellow contain blue cells and vice versa, as judged by comparing their fluorescence intensities with those of fully demixed blue and yellow sectors, see below. A second argument comes from the patch width of obligate mutualists remaining constant as the colony grows (Fig. 2D). Since the circumference increases during radial expansion, the number of patches at the circumference must increase to maintain a constant patch width. Indeed, we see new yellow patches emerge from within blue patches (Fig. 2C), which is only possible if the blue patch contain some yellow cells. Similarly, blue patches emerge from within yellow patches in Fig. 2C.

Incomplete demixing is due to frequency-dependent selection that promotes stable coexistence of two interacting strains [12, 14]. In our system, the selection coefficient

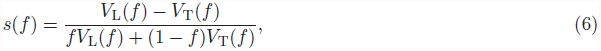

is frequency-dependent because the growth velocities *V*_L_ and *V*_T_ of the two strains depend on the amino acid concentrations *n*_L_ and *n_T_*, which in turn depend on the cellular fraction or ‘allele frequency’ *f* via Equation [2] and the corresponding tryptophan equation. Selection drives the system towards a stable fraction *f*^*^ with equal velocities, *V*_L_(*f*^*^) = *V*_T_(f^*^). Since our mutualistic interaction is symmetric, we predict *f*^*^ = 0.5.

**Figure 3:**
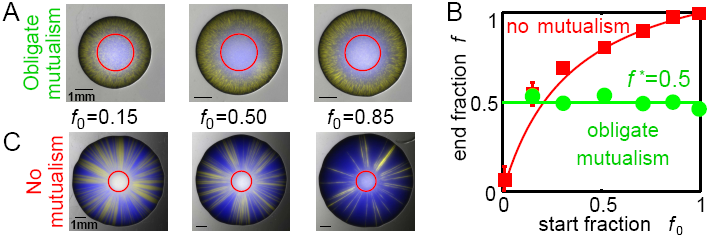
**A)** Obligate mutualists grow into a characteristic patchy pattern independent of the start fraction *f*_0_ in the homeland (red circle). **B)** The final fraction at the colony boundary equals *f* = *f*^*^ = 0.5 for obligate mutualism (green circles), but depends on the start fraction *f*_0_ for no mutualism (red squares). Solid lines are solutions to selection dynamics with selection coefficient *s*(*f*) of Equation [6] for obligate and *s* = 0.02 for no mutualism (blue cells are 2% more fit than yellow cells). **C)** Non-mutualistic colonies expand from the homeland (red circle) into demixed sectors whose number and width depends on the start fraction.

This prediction was confirmed experimentally: obligate mutualists inoculated with different starting fractions *f*_0_ expand into the same characteristic pattern with average fraction *f*^*^ = 0.5, independent of *f*_0_ (Fig. 3A,B). The transient dynamics towards steady-state growth depends on the start fraction: colonies with *f*_0_ = 0.15 are smaller because they take longer to start growing (SI Sec. S1.3), and colonies with *f*_0_ = 0.01 do not grow at all. While our simple model for steady-state growth captures the time scale of this transient, it does not capture these asymmetric features (SI Sec. S3.3).

For no mutualism, selection is not frequency-dependent, and the colony boundary fraction *f* depends on the start fraction *f*_0_ (Fig. 3B,C). The fraction *f* of blue cells increases during expansion because the blue tryptophan producers have a 2% fitness advantage over the yellow leucine producers under these conditions (Fig. 2A and SI Sec. S1.2), presumably because overproduction of tryptophan, a rare amino acid, is less costly than overproduction of the more abundant leucine.

### 2.3 Genetic drift can overcome facultative mutualism

That obligate mutualists remain (partially) mixed during spatial expansion is not completely unexpected since they *must* remain together to grow. We next study facultative mutualists, which can invade new territory on their own. Motivated by Fig. 2A, we decrease the mutualistic strength by increasing the amino acid concentrations in the medium. Fig. 4A shows the resulting colonies. For low leucine and tryptophan concentrations, expansion produces mixed patches with ragged boundaries, whereas large, well-separated sectors with smooth boundaries appear for high concentrations. When one amino acid is more abundant than the other, the strain requiring this amino acid has an advantage and dominates the colony. This behavior can also be seen in the colony boundary fraction *f* of blue cells (Fig. 4B). In summary, weak or asymmetric mutualism (high or asymmetric amino acid concentrations) leads to demixing, and strong mutualism (low concentrations) leads to mixing.

To probe the mutualism-drift antagonism, we focus on amino acid concentrations that retain the symmetry of our interaction, i.e. colonies whose boundary fractions *f* equal the inoculation fraction *f*_0_ = 0.5. These colonies (outlined in white in Fig. 4A) are slightly off the diagonal *n*_EL_ = *n*_ET_, presumably because of the slight fitness advantage of blue cells and the approximate nature of the linear fit in Fig. 2A.

To study the degree of mixing, we consider the local fraction *f* of blue cells at the colony boundary. In the demixed case, a histogram of *f* should have peaks at *f* = 1 (from blue sectors) and *f* = 0 (from yellow sectors), with only few 0 < *f* < 1 values (at sector boundaries). In contrast, more mixed mutualistic patches should exhibit a broad peak around *f* = 0.5. The yellow fluorescence intensity, a measure of the amount of yellow cells, displays this behavior (Fig. 4C): for low external amino acid concentrations, the histograms have a single peak, while they are clearly bimodal for higher concentrations. More quantitatively, as mutualism becomes weaker with increasing amino acid concentration *n*_E_ ≡ (*n*_EL_ + *n*_ET_)/2, the probability of seeing one peak in histograms of random subsets of the fluorescence intensity data drops sharply from 1 to 0 as *n*_E_ increases above 0.25 (Fig. 4D). Correspondingly, the patch width *L* increases with *n*_E_ until it plateaus for *n*_E_ ≳0.25 (Fig. 4E). We conclude that genetic drift overpowers mutualism for amino acid concentrations *n*_E_ ≳0.25, even though mutualism and complete mixing would still be beneficial, since *n*_E_ = 0.25 supports only 25% of the maximal growth rate of isolated, well-fed strains (Fig. 2A and Fig. 4F).

**Figure 4:**
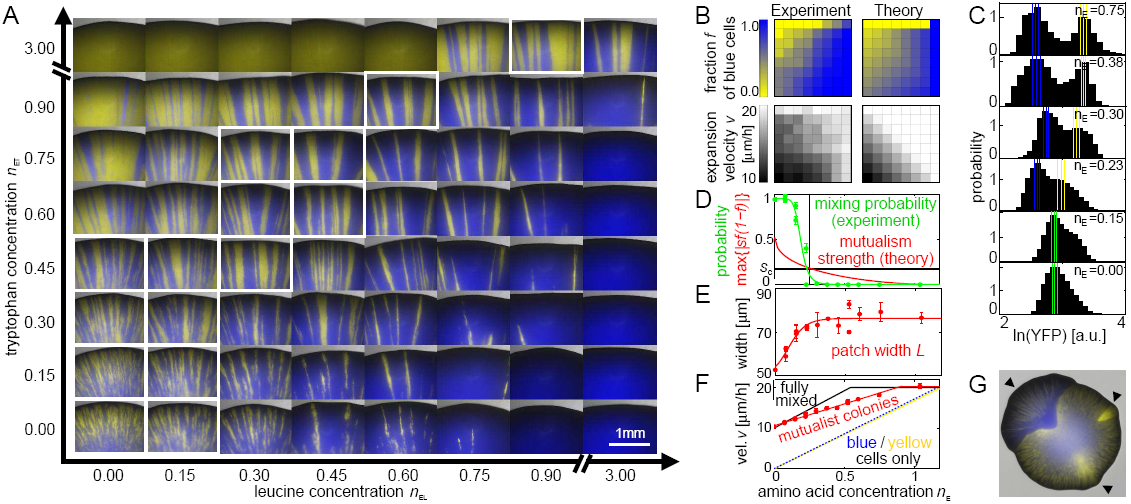
Tuning strength and symmetry of mutualism. **A)** Images of colony boundaries for different external leucine and tryptophan concentrations *n*_EL_ and *n*_ET_ in the medium. For low concentrations (lower left corner), colonies display a patchy pattern with ragged boundaries, while high external amino acid concentrations (upper right corner) lead to demixing into clear sectors with smooth boundaries. If leucine is abundant but tryptophan is not (lower right corner), the blue leucine-requiring strain wins, whereas abundant tryptophan and low leucine (upper left corner) favors the yellow tryptophan-requiring strain. Colonies with boundary fractions *f* within 10% of the inoculation fraction *f_0_* = 0.5 are outlined by white squares and further investigated in (C-F). **B)** Average colony boundary fraction *f* of blue cells (top) and expansion velocity *v* (bottom), as measured experimentally (left) and predicted by a model for well-mixed growth (right). The horizontal and vertical axes cover the same amino acid concentrations as in (A). **C)** Characterization of demixing along the ‘diagonal’ (white-square images in (A)). The yellow fluorescence intensity histograms display a single broad peak for small *n*_E_ = (*n*_EL_ + *n*_ET_)/2, and two peaks for large *n*_E_. Vertical lines indicate the mode locations (solid lines) and their standard deviations (dashed lines) for histograms from random data subsets. **D)** The probability of finding 1 peak (green points, green line to guide the eye) in fluorescence histograms of random data subsets decreases with increasing *n*_E_, exhibiting a sharp drop near *n*_E_ = 0.25 (vertical black line). The theoretically predicted strength of mutualism, estimated as the maximum of the selection term |*s*(*f*)*f* (1 – *f*)| of Equation [6] (red line), becomes comparable to the strength of genetic drift, *s_c_* = *D*^2^_g_/*D*_s_ (horizontal black line), at the concentration *n*_E_ = 0.25 of crossover from mixing to demixing. **E)** The average width *L* (red points, red line to guide the eye) of patches of a single color at distance *R*–*R*_0_ = 1.5 mm from the homeland correspondingly increases with *n*_E_ until it plateaus for *n*_E_ ≳ 0.25. **F)** The velocity of mutualistic colonies increases with *n*E (red data points), but not as fast as predicted by the mutualistic benefits (black solid line). However, it is always larger than the single strain growth velocities (yellow solid and blue dotted lines replotted from Fig. 2A). **G)** A colony of obligate mutualists with 3 mutant sectors indicated by black arrowheads. The average is one mutant sector per colony.

To understand how varying the mutualism’s strength controls the antagonism between mutualism and genetic drift, we write down a model for the dynamics of the cellular fraction *f* (*x*, *τ*) of blue cells [35, 36, 14]:

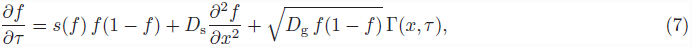

where *τ* is time measured in generations and *x* is the coordinate along the front. The first term incorporates mutualism with the selection coefficient *s*(*f*) of Equation [6]. The second term describes cellular diffusion due to the jostling of cell division with diffusion constant *D*_s_. The last term describes genetic drift, where Γ(*x*, τ) is an Itō delta-correlated Gaussian noise. The genetic diffusion constant *D*_g_ characterizes the noise strength and is expected to be inversely proportional to the effective population density at the front [35].

The selection term favors mixing as long as the amino acid concentrations are below the crossover values in Fig. 2A. In this regime, amino acids secreted by the cells increase growth velocities, so that mutualists benefit from remaining mixed at an optimal fraction *f*^*^. Considering only this term gives reasonable predictions for the average cell fraction *f* (top panels of Fig. 4B) as *f* depends mainly on the overall amino acid balance. However, it overestimates the growth velocities (compare the bottom panels in Fig. 4B, and the black solid line with the red data points in Fig. 4F). This discrepancy arises because genetic drift during spatial expansion leads to local deviations from the optimal fraction *f*^*^, thus slowing colony growth and producing blue and yellow patches instead of a homogeneous mix at fraction *f*^*^.

A detailed analysis of Equation [7], assuming locally flat fronts, shows that mutualism loses to genetic drift when the mutualistic strength falls below a critical value 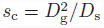 that characterizes the strength of local demixing [14]. Using our independently determined experimental parameters, the crossover concentration is predicted to be *n*_E_ = 0.25, which is consistent with our experimental observations (Fig. 4C–D). For asymmetric mutualism, e.g. due to more leucine than tryptophan in the medium, local fixation of leucine consumers (*f* =1) becomes more likely because the mutualistic fixed point *f*^*^ is closer to *f* = 1, and because the selective barrier to fixation at *f* = 1 is lower (SI Sec. S3 and Ref. [37]).

Obligate mutualists grow in a characteristic pattern determined by mutualism parameters such as nutrient uptake and secretion rates. On evolutionary time scales, mutations can change these properties and thus the pattern. To our surprise, mutant sectors arise even in our ~50 generation expansions (Fig. 4G). These mutants presumably change the mutualistic interaction since they expand with patterns different from the ancestors, and since most sectors have a different shape than expected for frequency-independent selection coefficients [38]. We plan to investigate these mutants further in the future.

## 3 Conclusions

During spatial population expansions, mutualism and genetic drift act as antagonistic evolutionary forces. Mutualists benefit from coexistence (‘mixing’). Due to this constraint, genetic drift at the expansion front can impede or even destroy mutualism by creating regions that are colonized predominantly or exclusively by one of the partners (‘demixing’). We experimentally and theoretically investigated this antagonism during the spatial expansion of two cross-feeding yeast strains. We find that strong mutualism suppresses demixing, but weaker mutualism is overpowered by genetic drift even though the resultant demixing makes the population grow more slowly than it would if it remained fully mixed. A critical mutualistic selection strength is required for mutualists to ‘survive’ spatial expansions. Our results are quantitatively explained by a model that incorporates mutualistic frequency-dependent selection due to nutrient exchange as well as the diffusional drift of cells due to cell division.

Spatial demixing is particularly pronounced in our experimental system because yeast lack motility and only ‘disperse’ offspring locally by cell division. For other organisms, movement of individuals and offspring dispersal provide additional mixing. But movement and dispersal are usually spatially restricted, and genetic demixing will occur if expansion into new territory is sufficiently fast compared to migration and dispersal within occupied areas [4, 34]. Spatial sectoring has been observed for mutualists in nature [39, 40].

The detrimental effect of spatial expansion on mutualism contrasts with the notion that spatial structure in general [41, 25] and spatial expansion in particular [8, 9, 10, 11] promote cooperation. In these studies, cooperators benefit from demixing, which separates them from non-cooperating ‘cheaters’. In contrast, mutualists profit from coexistence rather than separation and are impeded by expansions. The effect of spatial expansion, similar to spatial structure for stationary populations [42, 43], therefore depends on whether the cooperative interaction selects for a mixture of genotypes. If a mixture of genotypes grows fastest, spatial expansion impedes proliferation by separating these genotypes.

The difficulty of successfully expanding into new territories may contribute to mutualism breakdown [7], and explain the rareness in nature of mutualisms that form exclusively between two species [44]. Our observations suggest that mutualists can only expand their range together if mutualistic benefits are very strong, or if they can ensure coordinated dispersal of the mutualistic partners. Strong benefits presumably allowed the spread of flowering plants and their pollinators [17], the invasion of land by plants with mutualistic fungi [16], and underlie many current plant-microbe mutualisms [18, 19]. Other mutualisms exhibit permanent physical linkage, most notably eukaryotic cells and their mitochondria and chloroplasts, endosymbionts, and lichens. Indirect physical linkage can preserve mutualisms, such as agricultural ants transporting their fungal crops to new nests [45]. In summary, the requirement to ‘survive’ territorial expansions may have played an important role in the evolution of many mutualisms.

## 4 Materials and methods

**Strains** The haploid, asexually reproducing, *S. cerevisiae* strains Leu^FBR^ Trp^−^ and Leu^−^ Trp^FBR^ are derived from W303, can make all amino acids besides leucine and tryptophan, and share these markers: *MATa can*1-100 *hmlα*Δ::*BLE leu*9Δ::*KANMX*6 *prACT*1 – *yCerulean – tADH*1@*URA*3. Strain Leu^FBR^ Trp^−^ has the additional modifications *his*3Δ::*prACT*1 – *ymCitrine – tADH*1:*HIS*3*MX*6 *LEU*4^FBR^ *trp*2Δ::*NATMX*4, and strain Leu^−^ Trp^FBR^ has *his*3Δ::*prACT*1–*ymCherry–tADH*1:*HIS*3*MX*6 *leu*4Δ::*HPHMX*4 *TRP*2^FBR^. The enzymes Leu4^FBR^ (Ref.[31]) and Trp2^FBR^ (Ref. [32]) are insensitive to feedback-inhibition by leucine and tryptophan, respectively. Both strains express the cyan fluorescent protein yCerulean. Leu^FBR^ Trp^−^ also expresses the yellow fluorescent protein mCitrine, and Leu^−^ Trp^FBR^ the red fluorescent protein mCherry. To enhance contrast, Leu^FBR^ Trp^−^ and Leu^−^ Trp^FBR^ are depicted as yellow and blue, respectively. More strain are in the SI Sec. S1.

**Growth conditions** We used 1% agarose plates with CSM-leucine-tryptophan (complete synthetic medium as described in Ref. [46], except 2 mg/l of adenine and no leucine and tryptophan were used), plus appropriate amounts of leucine and tryptophan. For fully complemented CSM, we added at least 1524 *µ*M leucine and 196 *µ*M tryptophan. Cells were pre-grown in liquid CSM at 30°C in exponential phase for more than 12 h, counted with a Beckman Coulter counter, and mixed in appropriate ratios. The mix was spun down and vortexed after discarding the supernatant. A 0.5 *μ*l drop of the mix (≈ 10^9^ cells/ml) was pipetted on agar plates that had dried for 2 days post-pouring. Plates were incubated at 30°C in a humidified box for 7 days and imaged with a Zeiss Lumar stereoscope.

**Radial growth velocity** Colonies were imaged once a day, and their radii, determined by circle fitting with MATLAB, were fitted with a straight line for days 4-7. The velocity is the average of slopes from at least 3 different colonies.

**Boundary fraction** (*f*) Cells were scraped from colony boundaries with a pipet tip, avoiding mutant sectors, and resuspended in PBS. The fraction *f* of red fluorescent cells was determined on a Beckton-Dickinson LSR Fortessa flow cytometer.

**Patch width** (*L*) Using MATLAB, we determined the local maxima in the yellow fluorescence intensity (normalized by the cyan fluorescent intensity to correct effects of varying colony thickness and unequal lighting, and smoothed over 15 pixels) plotted along the circumference separately for each radius. The patch width *L* is the circumference divided by twice the number of maxima. Data within 20 pixels from the colony boundary were excluded because of weak fluorescence intensity. Mutant sectors were excluded from the analysis. Using the red fluorescence intensity instead of the yellow intensity gave similar results.

**Histograms to characterize demixing** We constructed the histogram of the yellow fluorescence intensity (normalized by cyan) of 3000 individual pixels that were randomly selected from the region at 50 to 550µm distance from the colony boundary, and determined the locations and number of its modes (separated by at least 3 bins). In a bootstrapping analysis, we performed this procedure 100 times to determine the average mode locations and the probability of observing only one mode. Using the red fluorescence intensity (normalized by cyan) gave similar results.

## Acknowledgements

We thank Erik F. Hom, Kirill S. Korolev, and J. David van Dyken for discussions. Support for this work was provided by the National Institute of General Medical Sciences Grant P50GM068763 of the National Centers for Systems Biology, by the National Science Foundation through grant DMR-1005289, and by the Harvard Materials Research Science and Engineering Center through grant DMR-0820484. MJIM was supported by a research fellowship from the German Research Foundation and a grant from the National Philanthropic Trust.

## References

[1] RM Donlan. Biofilms: Microbial life on surfaces. Emerg Infect Dis, 8:881–90, 2002.

[2] AR Templeton. Out of Africa again and again. Nature, 416:45–51, 2002.

[3] C Parmesan. Ecological and evolutionary responses to recent climate change. Annu Rev Ecol Evol Syst, 37:637–669, 2006.

[4] L Excoffier, M Foll, and RJ Petit. Genetic consequences of range expansions. Annu Rev Ecol Evol Syst, 40:481–501, 2009.

[5] O Hallatschek, P Hersen, S Ramanathan, and DR Nelson. Genetic drift at expanding frontiers promotes gene segregation. Proc Natl Acad Sci USA, 104:19926–30, 2007.

[6] A Hastings, K Cuddington, KF Davies, CJ Dugaw, S Elmendorf, A Freestone, S Harrison, M Holland, J Lambrinos, U Malvadkar, BA Melbourne, K Moore, C Taylor, and D Thomson. The spatial spread of invasions: new developments in theory and evidence. Ecol Lett, 8:91–101, 2005.

[7] ET Kiers, TM Palmer, AR Ives, JF Bruno, and JL Bronstein. Mutualisms in a changing world: an evolutionary perspective. Ecol Lett, 13:1459–1474, 2010.

[8] CD Nadell, KR Foster, and JB Xavier. Emergence of spatial structure in cell groups and the evolution of cooperation. PLoS Comp Biol, 6:e1000716, 2010.

[9] KS Korolev. The fate of cooperation during range expansions. PLoS Comp Biol, 9:e1002994, 2013.

[10] JD van Dyken, MJI Müller, KML Mack, and MM Desai. Spatial population expansion promotes the evolution of cooperation in an experimental Prisoners Dilemma. Curr Biol, 23:919–23, 2013.

[11] MS Datta, KS Korolev, I Cvijovic, C Dudley, and J Gore. Range expansion promotes cooperation in an experimental microbial metapopulation. Proc Natl Acad Sci USA, 110:7354–9, 2013.

[12] JN Holland and DL DeAngelis. A consumer-resource approach to the density-dependent population dynamics of mutualism. Ecology, 91:1286–1295, 2010.

[13] B Momeni, KA Brileya, MW Fields, and W Shou. Strong inter-population cooperation leads to partner intermixing in microbial communities. eLife, 2:e00230, 2013.

[14] KS Korolev and DR Nelson. Competition and cooperation in one-dimensional stepping-stone models. Phys Rev Lett, 107:088103, 2011.

[15] Luca Dall’Asta, Fabio Caccioli, and Deborah Beghe. Fixation-coexistence transition in spatial populations. EPL, 101, 2013.

[16] MA Selosse and F Le Tacon. The land flora: a phototroph-fungus partnership? Trends Ecol Evol, 13:15–20, 1998.

[17] WL Crepet. Progress in understanding angiosperm history, success, and relationships: Darwin’s abominably ‘perplexing phenomenon’. PNAS, 97:12939–41, 2000.

[18] DM Richardson, PA Williams, and RJ Hobbs. Pine invasions in the southern-hemisphere - determinants of spread and invadability. J Biogeogr, 21:511–527, 1994.

[19] DM Richardson, N Allsopp, CM d’Antonio, SJ Milton, and M Rejmánek. Plant invasions - the role of mutualisms. Biol Rev Camb Philos Soc, 75:65–93, 2000.

[20] SA West, SP Diggle, A Buckling, A Gardner, and AS Griffins. The social lives of microbes. Annu Rev Ecol Evol Syst, 38:53–77, 2007.

[21] RJ Palmer, K Kazmerzak, MC Hansen, and PE Kolenbrander. Mutualism versus independence: Strategies of mixed-species oral biofilms in vitro using saliva as the sole nutrient source. Infect Immun, 69:5794–5804, 2001.

[22] T Pfeiffer and S Bonhoeffer. Evolution of cross-feeding in microbial populations. Am Nat, 163:E126–E135, 2004.

[23] W Shou, S Ram, and JMG. Vilar. Synthetic cooperation in engineered yeast populations. Proc Natl Acad Sci USA, 104:1877–1882, 2007.

[24] HJ Kim, JQ Boedicker, JW Choi, and RF Ismagilov. Defined spatial structure stabilizes a synthetic multispecies bacterial community. Proc Natl Acad Sci USA, 105:18188–18193, 2008.

[25] W Harcombe. Novel cooperation experimentally evolved between species. Evolution, 64:2166–2172, 2010.

[26] KL Hillesland and DA Stahl. Rapid evolution of stability and productivity at the origin of a microbial mutualism. Proc Natl Acad Sci USA, 107:2124–2129, 2010.

[27] EH Wintermute and PA Silver. Emergent cooperation in microbial metabolism. Mol Sys Biol, 6:407, 2010.

[28] SJ Pirt. A kinetic study of mode of growth of surface colonies of bacteria and fungi. J Gen Microbiol, 47:181–197, 1967.

[29] MO Lavrentovich, JH Koschwanez, and DR Nelson. Nutrient shielding in clusters of cells. Phys Rev E, 87:062703, 2013.

[30] O Hallatschek and DR Nelson. Gene surfing in expanding populations. Theor Popul Biol, 73:158–170, 2008.

[31] D Cavalieri, E Casalone, B Bendoni, G Fia, M Polsinelli, and C Barberio. Trifluoroleucine resistance and regulation of alpha-isopropyl malate synthase in *Saccharomyces cerevisiae*. Mol Gen Genetics, 261:152–160, 1999.

[32] R Graf, B Mehmann, and GH Braus. Analysis of feedback-resistant anthranilate synthases from *Saccharomyces cerevisiae*. J Bacteriol, 175:1061–1068, 1993.

[33] H Akashi. Translational selection and yeast proteome evolution. Genetics, 164:1291–1303, 2003.

[34] O Hallatschek and DR Nelson. Life at the front of an expanding population. Evolution, 64:193–206, 2009.

[35] KS Korolev, M Avlund, O Hallatschek, and DR Nelson. Genetic demixing and evolution in linear stepping stone models. Rev Mod Phys., 82:1691–1718, 2009.

[36] E Frey. Evolutionary game theory: Theoretical concepts and applications to microbial communities. Physica A, 389:4265–4298, 2010.

[37] MO Lavrentovich and DR Nelson. Asymmetric mutualism in two and three dimensions. arXiv:1309.0273v1, 2013.

[38] KS Korolev, MJI Müller, N Karahan, AW Murray, O Hallatschek, and DR Nelson. Selective sweeps in growing microbial colonies. Phys Biol, 9:026008, 2012.

[39] AS Mikheyev, T Vo, and UG Mueller. Phylogeography of post-pleistocene population expansion in a fungus-gardening ant and its microbial mutualists. Mol Ecol, 17:4480–4488, 2008.

[40] DM Althoff, KA Segraves, CI Smith, J Leebens-Mack, and O Pellmyr. Geographic isolation trumps coevolution as a driver of yucca and yucca moth diversification. Mol Phylogenet Evol, 62:898–906, 2012.

[41] MA Nowak. Five rules for the evolution of cooperation. Science, 314:1560–3, 2006.

[42] CP Roca, JA Cuesta, and A Sanchez. Effect of spatial structure on the evolution of cooperation. Phys Rev E, 80:046106, 2009.

[43] DB Borenstein, Y Meir, JW Shaevitz, and NS Wingreen. Non-Local Interaction via Diffusible Resource Prevents Coexistence of Cooperators and Cheaters in a Lattice Model. PLoS One, 8:e63304, 2013.

[44] HF Howe. Constraints on the evolution of mutualisms. Amer Nat, 123:764–777, 1984.

[45] UG Mueller, NM Gerardo, DK Aanen, DL Six, and TR Schultz. The evolution of agriculture in insects. Annu Rev Ecol Syst, 36:563–595, 2005.

[46] D Burke, D Dawson, and T Stearns. Methods in Yeast Genetics. Cold Spring Harbor Laboratory Press, USA, 2000.

